# An auditable harness for biomolecular dynamics simulations by AI agents

**DOI:** 10.64898/2026.11.09.765490

**Authors:** Yasuhiro Matsunaga, Yichao Wu, Yangyang Zhang, Shweta Kumari, Akari Enomoto, Yudai Homma, Shin Hatakeyama, Ai Shinobu

## Abstract

AI agents now generate biological hypotheses faster than they can be tested. Hypotheses stated at the atomic level can be examined with molecular dynamics (MD) simulations. For new questions without a reference, trust in an agent-run simulation rests on a tool-written process record, not the agent’s account. Existing MD agents leave this record to the language model. Here we present MDClaw, a harness whose validating tools and directed acyclic graph (DAG) check stages before execution and record every simulation stage. On MDDataBench, which grades outputs against deposited simulations, four open-weight models with MDClaw passed all three runs on 81–100% of 31 test systems. The DAG complements this outcome check. An ablation without the DAG revealed 11 successful runs that had skipped minimization or equilibration, or simulated a different structure. MDClaw carried questions from human and AI scientists through simulation to trajectories traceable to their inputs and settings.

---

In biological research, AI systems increasingly help scientists synthesize literature, formulate hypotheses and design studies (1). Robin and Co-Scientist generate and prioritize research proposals (2, 3). Biomni combines biomedical tools to execute analyses from natural-language requests (4). Domain-specific agents analyze biological data and orchestrate chemistry tools (5, 6). Coding agents can also carry out substantial portions of computational research (7, 8). These proposals become evidence only when they are tested by experiments or simulations that others can examine.

Molecular dynamics (MD) simulations can test hypotheses stated at the atomic level, such as how a mutation destabilizes a fold or how a ligand engages its binding site. Mature simulation engines provide the numerical machinery (9, 10). The difficult part is translating a scientific question into a molecular system and a simulation protocol. A membrane channel, a metalloprotein and a protein–ligand complex each need different choices of composition, protonation, forcefield parameters and solvent. Each choice affects every later stage, from minimization to analysis. Workflows such as CHARMM-GUI make these choices reliable for human users by encoding expert protocols (11).

When an AI agent runs the simulation, its result can be compared with a reference only for benchmark tasks. For a new research question, no reference exists, and whether to trust the result depends on how it was obtained: which calculations were run, from which inputs and under which conditions. Even when a reference exists, comparing final files misses errors in intermediate stages. An agent can skip equilibration or start production from the wrong coordinates and still produce files that pass. The process therefore has to be checkable. This requires a record that the tools write as they run, not the agent. The agent’s own summary cannot serve this purpose, because it can sound correct even when the work behind it was not. Such a record lets others reconstruct what an autonomous system did, audit it and correct it when it is wrong (12).

Existing MD agents show that language models can set up and run simulations, but they leave the record of the process to the model. MDCrow provides expert-designed simulation and analysis tools (13), MDAgent combines planning and supervision with reusable case knowledge (14), and MD-Forge generates pipelines and revises them from simulator feedback (15). MutexaGPT translates enzyme-engineering requests into physical models and design proposals (16). Related agents address materials and computational-chemistry simulations (17–19) and protein design (20, 21), and MD-LLM-1 applies a language model directly to molecular dynamics (22). In most of these systems, the language model decides the order of stages and passes files and conditions from one stage to the next. The record of what was run is mainly the agent’s own log and summary. This record depends on the same agent whose work it is meant to check.

A harness also has to be tested on tasks where the correct outcome is known. This requires reference data, which is hard to obtain for MD. A simulation has no single correct trajectory, and references generated with the benchmark authors’ own workflow grade agents against that workflow’s choices. Existing benchmarks therefore use task-specific criteria. MDArena combines output checks with a language-model judge of execution logs across 50 tasks (23), and MDGym supplies starting structures and force-field files and scores computed properties within a 5% tolerance (24). Benchmarks in other scientific domains reveal sub-stantial gaps (25, 26). MDDataBench adds a comparison of an agent’s prepared system with an independent reference topology, and of its dynamics with deposited simulation data. The MDDB federation makes such references available at scale. Researchers have deposited tens of thousands of simulation datasets, each grouping trajectories that share a topology (27), as part of efforts to make simulation data findable, accessible, interoperable and reusable (28). An agent’s output can therefore be compared with simulations that neither the agent nor the benchmark authors produced.

Here we present MDClaw, a harness for coding agents, that is, the tools, instructions and records that surround the language model. MDClaw combines protocol documents (“agent skills”), validating command-line tools and a persistent directed acyclic graph (DAG) of calculation stages. Because an agent builds its workflow while it runs, the record cannot be designed in advance. Instead, each tool records its calculation as a node in the DAG and runs only with inputs from recorded earlier stages, so a file produced outside the record cannot enter a later stage. The same record lets errors be corrected without starting over. A researcher or agent can trace a problem back to the stage where it began and branch a new calculation from the last valid stage. We also introduce MDDataBench, which compares prepared systems and short trajectories with MDDB reference simulations. On MD-DataBench, MDClaw tools without skills already account for most of the gain in repeated success over the simulation software alone, and skills add a smaller increase. The DAG complements this outcome check by recording the process. An ablation that removed the DAG revealed 11 successful runs that had skipped minimization or equilibration, or simulated a different structure. Finally, MDClaw carries research questions from human scientists on Abl1 and hypotheses from the AI scientist Biomni on PNG-1 through simulation to results. No reference exists for these questions, and each trajectory can be checked by tracing its stages back to their inputs and settings.

## Results

### A DAG constrains execution and preserves the evidence behind a result

MDClaw connects a scientist’s hypothesis or prompt to a coding agent that plans simulations, executes them and analyzes the results (Fig. 1a). The scientist can be a human researcher or an AI system. An expert-designed harness provides 14 protocol documents, termed agent skills, and 100 command-line tools spanning structure preparation through trajectory analysis (Fig. 1b). Skills guide planning, while tools perform the calculations and enforce technical checks. The agent can select and revise a workflow while the harness retains its inputs, settings and outputs. The tools also run enhanced-sampling and alchemical free-energy methods (Supplementary Fig. 2).

**Fig. 1.**
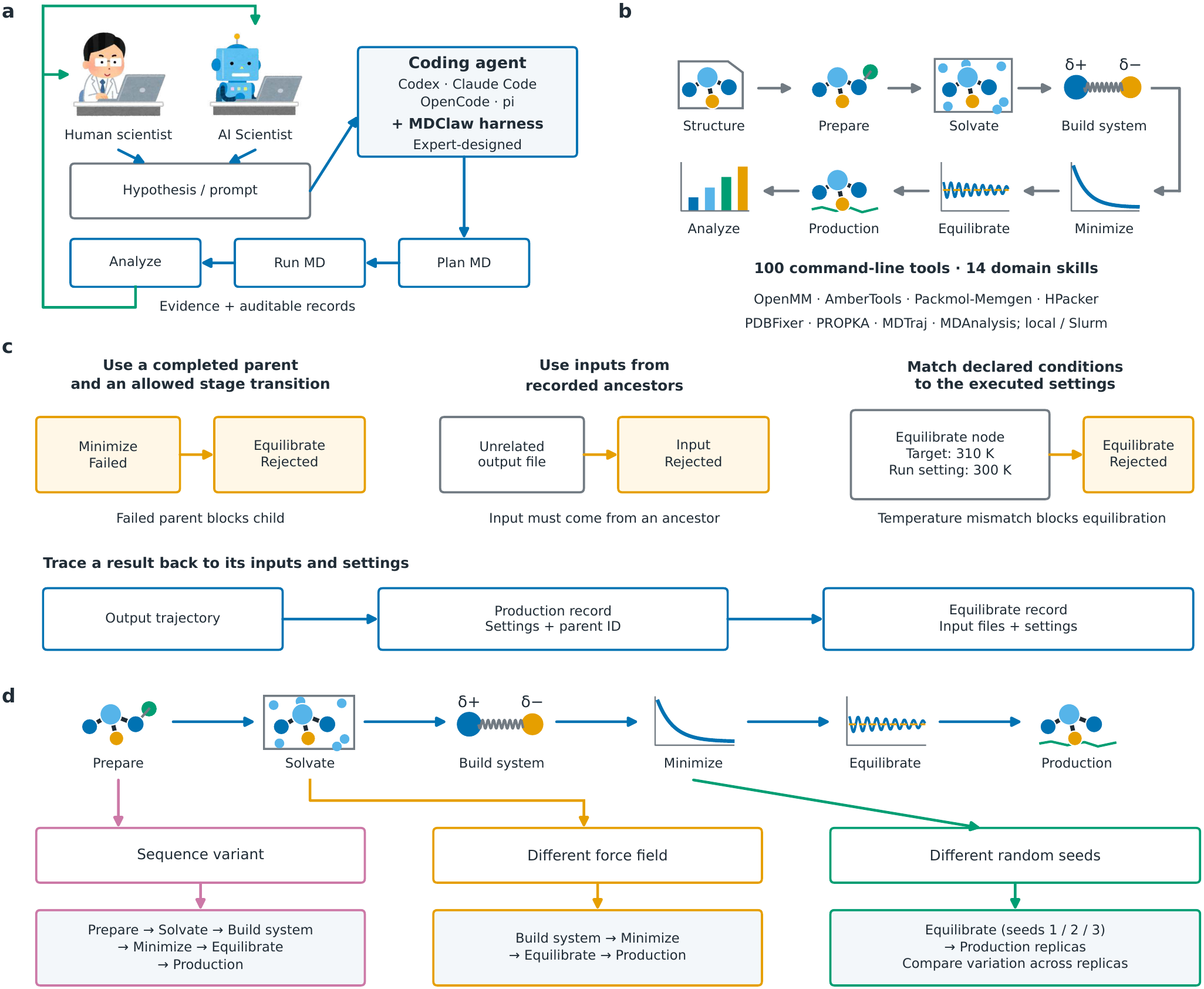
MDClaw uses an execution graph to constrain tool calls, retain auditable records and support branching workflows. **a**, A human scientist or AI Scientist supplies a hypothesis or scientific prompt. A coding agent, such as Codex, Claude Code, OpenCode or pi, uses the expert-designed MDClaw harness to plan and run MD calculations, analyze their results, and return evidence relevant to the hypothesis together with auditable records. **b**, MDClaw provides 100 command-line tools and 14 domain skills. The illustrated tools cover structure preparation, simulation and analysis. Build system includes force-field assignment. Arrows follow the illustrated workflow across the top row and back along the lower row. Execution uses OpenMM, AmberTools, Packmol-Memgen, HPacker, PDBFixer, PROPKA, MDTraj and MDAnalysis locally or through Slurm. **c**, General DAG rules and illustrative refusals. Executing a child requires a completed parent and an allowed stage transition (failed *Minimize* prevents *Equilibrate*). Inputs must follow recorded ancestry (an unrelated file is rejected). Declared conditions must match execution settings (an Equilibrate node targeting 310 K rejects a run configured at 300 K). The lower example traces an output trajectory to its Production record, including settings and the parent identifier, and then to the parent Equilibrate record and its input files. This allows results to be audited independently of the agent’s account. These rules reject the specified inconsistencies. They do not guarantee that all errors are prevented. **d**, Domain skills guide planning and command-line tools execute the plan. An agent can extend a study by reusing recorded inputs and choosing where to branch. Sequence variants restart at *Prepare*. Compatible force-field changes reuse *Solvate* outputs and restart at *Build system*. Replica calculations branch from the same completed *Minimize* node into *Equilibrate* with different random seeds, followed by *Production*. Comparing the replicas assesses run-to-run variation. Changing seeds does not by itself establish statistical independence or convergence. Stage names and illustrations match b.

The central data structure is a directed acyclic graph (DAG), in which nodes represent calculation stages and edges record their dependencies. The DAG also constrains tool calls (Fig. 1c): a stage can execute only when its required parent is complete, the transition is allowed, inputs belong to the recorded ancestry, and execution settings agree with declared conditions. For example, a failed Minimize node cannot supply an executable Equilibrate stage, and a node declared at 310 K cannot execute with a 300 K configuration. Chemical and force-field checks reject specified inconsistencies before expensive calculations begin. The tool returns a structured error to the agent that states why the call was rejected and suggests a corrective action, so the agent can branch from the last valid stage instead of starting over (Fig. 4c,d).

Each node retains parameters, file references, status and parent identifiers. A trajectory can be traced to the preparation and equilibration that produced it, independently of the agent’s summary. An agent can plan extensions from appropriate ancestors: mutations require a new preparation branch, compatible force-field changes can reuse solvation, and repeated calculations can branch into separate equilibrations with different random seeds (Fig. 1d). This preserves earlier calculations while allowing the research plan to evolve. Tools can assemble draft Methods, software citations and MDDB deposition bundles from the recorded workflow (Supplementary Fig. 1).

### MDDataBench evaluates submitted files against reference simulations

We developed MDDataBench to assess whether agents produce the requested molecular systems and short MD trajectories (Fig. 2). A survey of eight repositories in the Molecular Dynamics Data Bank (MDDB) identified 35,602 deposited simulation datasets. An initial eligibility screen across the three largest nodes yielded 4,036 entries with suitable topologies, trajectories, analyses and licences. We selected 128 development tasks spanning nine system classes. A separate test set contains 31 systems across eight classes, including complexes, nucleic acids and a membrane protein (Fig. 2a; Supplementary Table 1). To keep the test systems distinct from those MDClaw was developed on, no protein or nucleic-acid chain in a test system was similar in sequence to a development system or to a structure used in the MDClaw code (Methods).

**Fig. 2.**
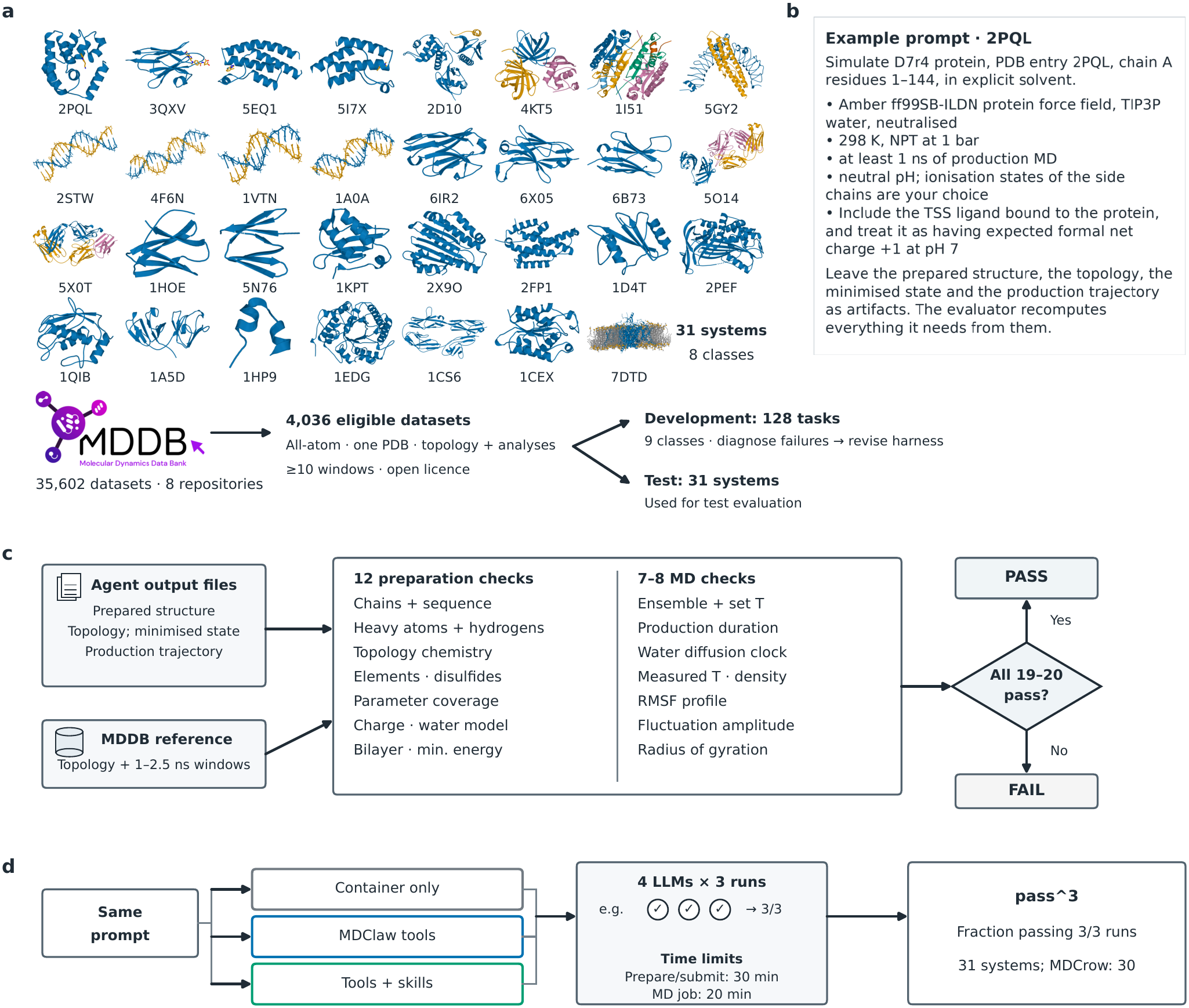
MDDataBench task systems, prompts and artifact-based scoring. **a**, All 31 test systems in task order, shown from their deposited structures. 7DTD includes the reference DPPC bilayer, with phosphate groups shown in orange and lipid tails in grey. Foreground lipids are omitted from the view to expose the protein. Structures use independent display scales and are not agent-generated results. The curation flow separates the 128 development tasks from the 31 test tasks. **b**, Image of the actual 2PQL prompt, with its boundary conditions and required artifacts. **c**, Scoring flow from submitted files and reference data to twelve preparation checks and seven or eight scored MD checks. Structural dynamics use reference windows of 1–2.5 ns. A pass requires all scored checks: 20 for 30 systems and 19 for 5X0T, where radius of gyration is diagnostic only (Methods). Agent reports are not scored. **d**, The same prompt is evaluated under three conditions, with repeated runs aggregated into the primary metric, `pass^3`, the fraction of systems for which all three runs pass. For these three main conditions, four large language models (LLMs) each perform three independent runs per system and condition, with time limits of 30 minutes for preparation/submission and 20 minutes per MD job. A separate MDCrow comparison uses two LLMs and 30 non-membrane systems, with its execution settings described in Methods.

A script generates each prompt from the metadata and files of the deposited entry, stating the target structure, molecular composition, force field, solvent, thermodynamic conditions and required outputs (Fig. 2b). The grader evaluates files rather than the agent’s account: twelve checks assess preparation and eight assess MD (Fig. 2c). Fluctuation profiles, fluctuation amplitude and radius of gyration are compared with tolerances calibrated from reference 1–2.5 ns windows. All scored checks must pass: normally 20, with 19 for 5X0T, whose radius-of-gyration check is diagnostic only (Methods). This measures conformity to a defined simulation task, rather than equilibrium convergence or biological validity.

We ran Kimi-K3, DeepSeek-V4.1-flash, GLM-5.3-flash and Qwen3.8-27B in the pi coding agent under three conditions: the simulation software container alone, MDClaw tools, and MDClaw tools plus skills. Each model ran the same prompt three times per system and condition (Fig. 2d). The primary metric, pass^3, is the fraction of systems passing all three runs. The pass@1 metric is the fraction of successful individual runs. Development tasks guided improvement, and the test tasks evaluated the pre-optimization and optimized harness versions.

### Large-scale development runs guide harness optimization

We developed the harness in repeated rounds on the 128 development tasks (Fig. 3a). After each round, coding agents inspected failed DAG nodes, inputs and execution records to diagnose errors, an MD specialist judged the proposed remedies, and we revised the tools or skills before the next round. We changed only the harness, not the language-model weights. Over these rounds we ran at least 3,726 agent runs on the RIKYU supercomputer at RIKEN R-CCS, across multiple language models and three tool-and-skill conditions, with three repeats per evaluated combination. The count includes unsuccessful attempts, not only completed trajectories. To assess the effect of this development on new systems, we compared the pre-optimization and optimized versions on the same 31 test tasks (Fig. 3b).

**Fig. 3.**
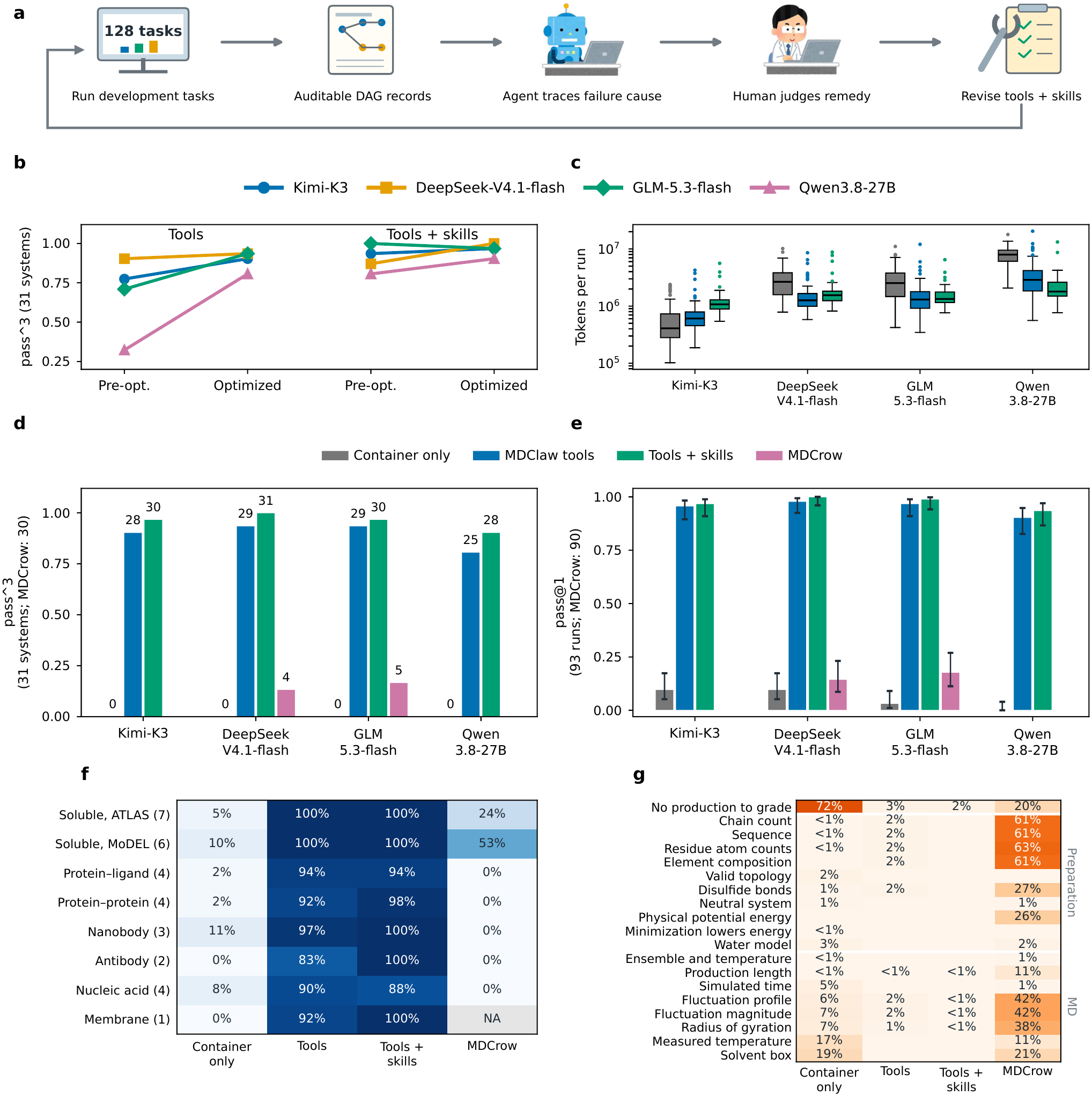
Harness optimization, performance and token usage on 31 MDDataBench test tasks. **a**, Repeated development rounds: an agent traces failed steps and their inputs through auditable DAG records to identify failure causes, a human specialist judges the remedy, and tools and skills are revised before the next round. The loop uses 128 development tasks. Language-model weights are unchanged. **b**, Pre-optimization and optimized harness versions evaluated on the same 31 test systems, with tools alone and tools plus skills. Each line represents one language model. **c**, Recorded prompt and output token usage per run for the three main conditions, using the condition colours shown above d and e and excluding MDCrow (*n* = 93 per group). All 1,116 included runs have complete token records. Reasoning tokens are included within output and not counted twice. Boxes show medians and quartiles. Whiskers extend to the most distant observations within 1.5 interquartile ranges. Points beyond the whiskers are shown individually. Median token usage is lower with MDClaw tools, with or without skills, than with the container alone for DeepSeek, GLM and Qwen. Kimi-K3 shows the opposite pattern. **d**, Fraction of systems passing all three runs (`pass^3`). Labels give system counts. Models are ordered Kimi-K3, DeepSeek-V4.1-flash, GLM-5.3-flash and Qwen3.8-27B. MDCrow was evaluated only with DeepSeek and GLM. Its denominator is 30 non-membrane systems, versus 31 systems for the other conditions. **e**, Single-run success (pass@1, *n* = 93 runs per bar, or *n* = 90 for MDCrow), with error bars showing 95% Wilson confidence intervals. The unsupported membrane task is excluded from MDCrow aggregates in d and e. The comparison therefore uses different task scopes. **f**, Percentage of successful runs by system class, pooled over four models for the main conditions and two for MDCrow. Darker blue indicates higher success. Percentages are printed in each cell and parentheses give system counts. MDCrow does not support membrane systems, so that cell is marked NA. **g**, Grader checks failed in unsuccessful runs, as a percentage of all runs of each condition (optimized version; MDCrow on the 30 non-membrane systems). A run without a gradable production is counted once in the top row. For the other unsuccessful runs each failed check is counted, so one run can appear in several rows. Only checks that failed at least once are shown; empty cells are 0%.

Of 124 model–task pairs (four models, 31 tasks), the number passing all three runs rose from 84 to 111 with tools alone and from 112 to 119 with tools plus skills. Tools alone gained the most, so the gap between the two conditions narrowed from 28 to eight pairs, consistent with procedural knowledge moving from the skills into checks and error messages in the tools. This shift is desirable because checks in the tools use none of the agent’s context and apply even without a skill, so the agent can choose its own route flexibly while the tools still enforce the DAG. This comparison is between complete releases, including container dependencies, and does not isolate individual changes.

### The harness improves repeated success

Without MD-Claw, 21 of 372 individual runs passed (pass@1 5.6%, 95% Wilson confidence interval 3.7–8.5%). With tools alone, pass^3 reached 25–29 of 31 systems (81–94%) and pass@1 reached 90–98% across models (Fig. 3d,e). Adding skills increased these ranges to 28–31 systems (90–100%) and 94–100%, respectively. Most of the improvement was already present with the command-line harness, and skills added to it. For DeepSeek, GLM and Qwen, MDClaw cut median token usage by 42–77%, as validated commands replaced trial-and-error scripting and model calls fell. Kimi-K3 was the exception, because without MDClaw it submitted its scripts after few model calls and 84 of 93 such runs failed (Fig. 3c).

We also evaluated MDCrow (13), a dedicated LLM agent for MD with its own built-in tools, whereas MDClaw equips general-purpose coding agents. It ran with DeepSeek and GLM, using the same prompts and grader. On the 30 non-membrane systems, MDCrow passed all three runs on four and five systems, respectively. The pass@1 rates were 13/90 and 16/90. All successes were soluble single-chain targets.

MDCrow does not support membrane proteins, so the membrane task was excluded from its evaluation (Fig. 3f). With MDClaw, every run on soluble proteins passed, and the other system classes passed 83–100% of runs.

Figure 3g shows which grader checks failed in each condition. Without MDClaw, 72% of runs ended without a production trajectory to grade, and graded runs failed mainly on the solvent box (19% of runs) and the measured temperature (17%). MDCrow usually produced a trajectory, but of a different system: 61–63% of its runs failed the chain count, sequence and composition checks, and 38–42% failed the fluctuation and radius-of-gyration checks. With MDClaw, 2–3% of runs had nothing to grade and no single check failed in more than 3% of runs.

### Removing the DAG reveals process errors in successful runs

To separate the contribution of the DAG, we built a version of MDClaw without it and evaluated it on the 30 non-membrane test systems with the same four models and prompts, three runs per system and tools without skills (Methods). The chemistry code and validating checks were unchanged. Each tool instead reads files from paths the agent supplies and writes to a directory the agent names, and no settings are inherited from earlier stages, so the agent carries every file and condition from one stage to the next. Without the DAG, pass^3 was 103 of 120 model–system pairs, compared with 108 with the DAG (two-sided sign test, *P ≈*0.30). This took more failed attempts: tool failures rose from 41 to 147 episodes and failed MD jobs from 1 to 23.

The DAG complements this outcome check by recording the process. Without the DAG, each tool call wrote a record of its input and output files with content hashes, from which we reconstructed the stages behind each submitted trajectory (Methods). Among 334 successful runs without the DAG, 11 had errors in the process (Fig. 4a). In six, equilibration started from the output of the topology build rather than from minimization. In two, production did as well, so neither minimization nor equilibration preceded the scored trajectory (Fig. 4b). In three, the simulated structure was not the prepared one, because the agent had changed or swapped it with its own files. With the DAG, none of the 343 successful runs had such errors. All 11 runs passed every output check, which compares the results with the reference, not the steps behind them. Without a reference, as in research, only a record of the process such as the DAG can reveal such errors.

**Fig. 4.**
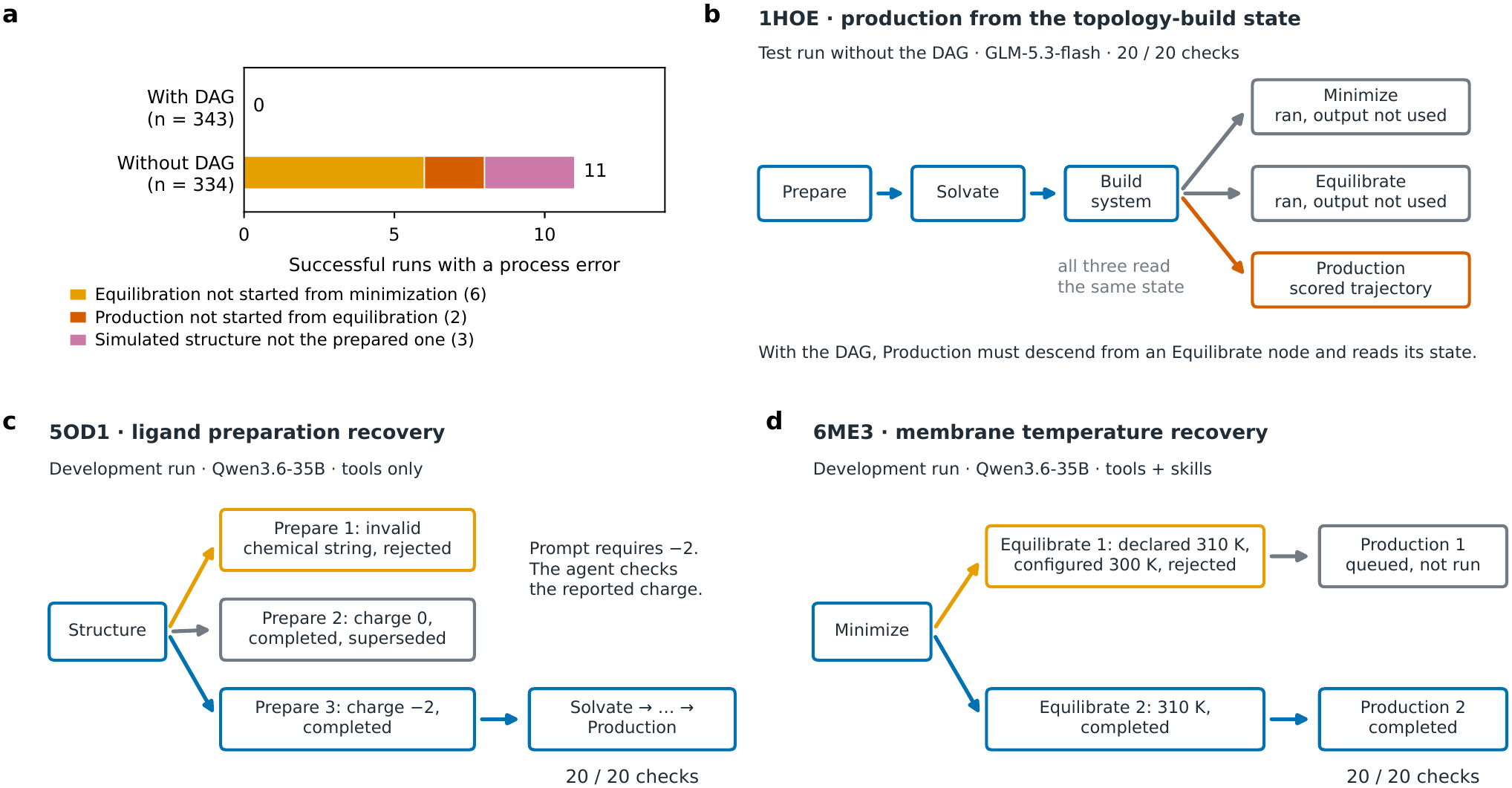
Removing the DAG reveals process errors in successful runs, and DAG records support recovery. **a**, Successful test runs with a process error, with and without the DAG (30 non-membrane systems, four models, three runs per system, tools without skills). With the DAG, the lineage of each scored production was read from the DAG records. Without the DAG, it was reconstructed from per-call records of input and output files with content hashes. *n* is the number of successful runs. Colours give the type of error, with counts in parentheses. **b**, A successful run without the DAG (1HOE, GLM-5.3-flash). Minimization, equilibration and production all read the state written by the topology build, so the scored trajectory did not start from the output of minimization or equilibration. With the DAG, a production node must descend from an equilibration node and reads its state. **c**, Ligand recovery in a development task (5OD1, Qwen3.6-35B, tools only). The tool rejects an invalid chemical string in Prepare 1. Prepare 2 completes but reports charge zero. The agent compares this with the prompt’s required charge of *−*2 and creates Prepare 3 from the same structure, which continues to production and passes all 20 checks. Prepare 2 remains recorded as completed. **d**, Temperature recovery in a development membrane task (6ME3, Qwen3.6-35B, tools plus skills). Equilibrate 1 is rejected before simulation because its configured 300 K conflicts with the declared 310 K, and its queued Production 1 is not run. Equilibrate 2 branches from the same Minimize node at 310 K and leads to Production 2, which passes all 20 checks. The development examples in c and d are separate from the test evaluation.

### Recorded branches support recovery from rejected calculations

Retained records also let agents recover from rejected calculations without repeating earlier stages (Fig. 4c,d). In a development ligand task (5OD1), the tool rejected an invalid chemical string. A second preparation completed but reported a ligand charge of zero, which the agent found to conflict with the prompt’s charge of *−*2. It created a third preparation from the same structure, which passed all 20 checks. In a development membrane task (6ME3), an Equilibrate node configured at 300 K, rather than the declared 310 K, was rejected before simulation. The agent branched a new Equilibrate node from the completed Minimize node at 310 K and passed all 20 checks.

### Simulations address human scientists’ research questions

We next used MDClaw to address two questions about Abl1 substrate recognition and peptide-scaffold engineering (Fig. 5a,b), realizing the human scientist–agent interaction outlined in Fig. 1a. The scientists posed the questions and designed the comparisons. They checked the selected models and fixed shared conditions. Agents carried out the rest with MDClaw: Boltz-2 co-folding and screening, system building, equilibration and production, all recorded in the DAG. Analyses ran outside the DAG, and the cyclic scaffold required four patches to MDClaw.

**Fig. 5.**
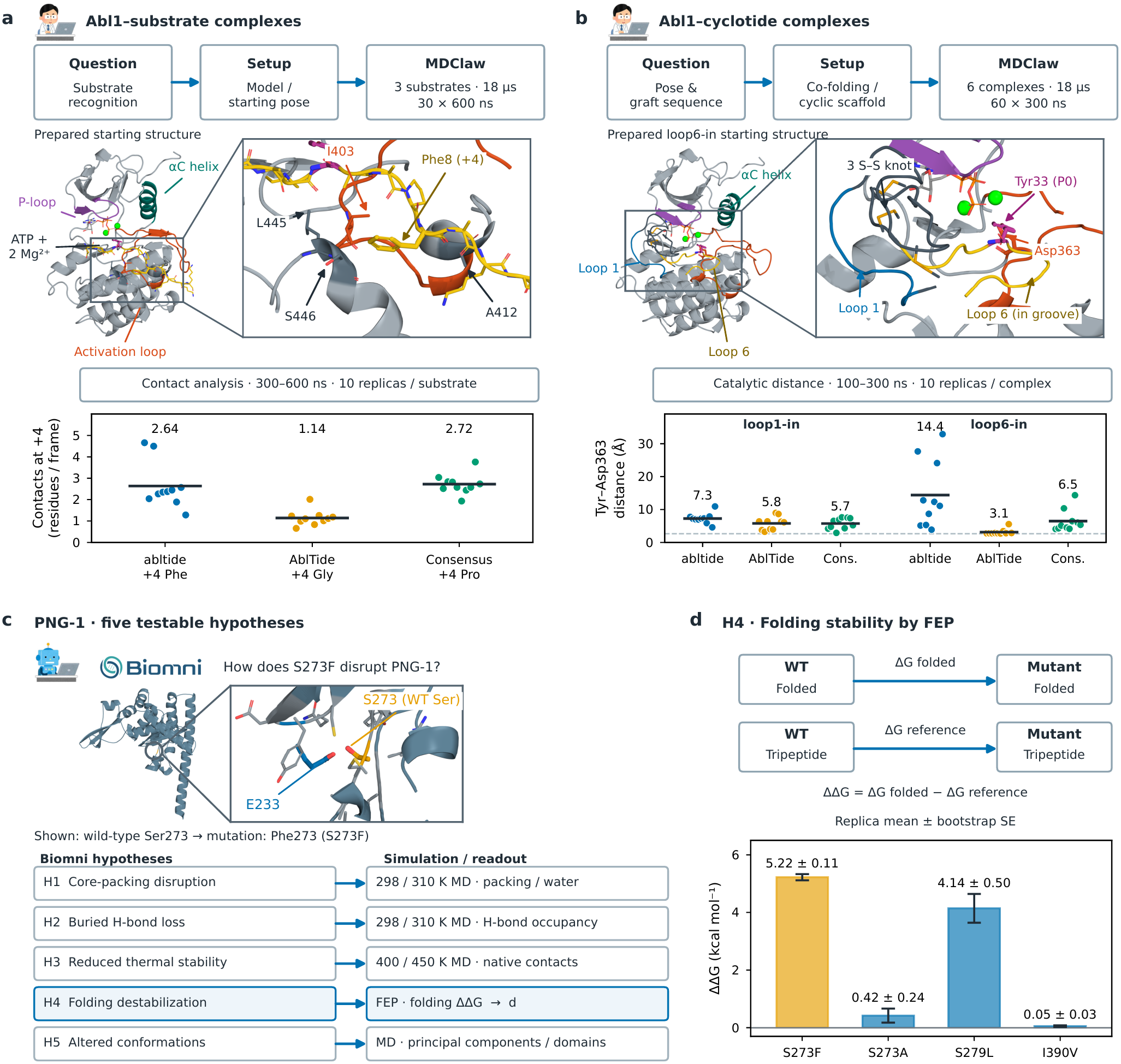
MDClaw connects human scientists’ research questions and an AI scientist’s hypotheses to simulations. **a**,**b**, Human scientists’ research requests, setup choices and MDClaw calculations in the linear Abl1–substrate and Abl1–cyclotide studies. The human and AI scientist icons in a–c link these applications to Fig. 1a. Points show trajectory averages and horizontal marks their arithmetic means. The molecular illustrations provide structural context. **a**, Prepared Abl1–abltide structure before minimization and equilibration, with Phe8 (+4) and nearby kinase residues enlarged. The peptide is yellow, activation loop orange, P-loop purple, *α*C helix teal and Tyr4 (position 0) magenta. Ten 600 ns replicas per substrate totalled 18 *µ*s. The plot shows contacting kinase residues per frame over 300–600 ns (4.5 Å heavy-atom cutoff). Means are 2.64, 1.14 and 2.72. This checks the trend in the earlier 150–300 ns interval of the same trajectories. **b**, Prepared loop6-in Abl1–cyclotide starting structure before minimization and equilibration, with acceptor Tyr33 (magenta), catalytic base Asp363 (orange), in-groove loop 6 (yellow), loop 1 (blue) and three cystine-knot disulfides enlarged. Six complexes (two binding poses *×* three in-groove sequences, with the non-engaged loop held as Abl Consensus) were simulated for ten 300 ns replicas each, totalling 18 *µ*s. The plot shows replica-mean distances from the in-groove acceptor tyrosine OH (Tyr5 in loop1-in, Tyr33 in loop6-in) to the Asp363 carboxylate over bound frames in 100–300 ns. The dashed line marks the 2.65 Å crystallographic reference (PDB 1IR3). Means are 7.3, 5.8 and 5.7 Å in loop1-in, and 14.4, 3.1 and 6.5 Å in loop6-in. **c**, Five Biomni hypotheses about S273F in PNG-1, a *Neurospora crassa* peptide:*N*-glycanase orthologue required for cell polarity, and their corresponding simulation/readout plans: core-packing disruption (H1), buried hydrogen-bond loss (H2), reduced thermal stability (H3), folding destabilization (H4) and altered conformations (H5). The structure locates the S273F substitution site in the initial wild-type model. The enlarged region shows the wild-type Ser273 side chain (orange), nearby Glu233 (blue) and surrounding side chains as sticks. It does not depict a phenylalanine mutant or an observed mutation-induced change. The H4 row links to the FEP result in d. **d**, FEP evaluation of the folding-stability hypothesis H4. Alchemical transformations in the folded construct and a tripeptide reference give ΔΔ*G* = Δ*G*_folded_ *−* Δ*G*_reference_. The tripeptide is a reference model, not a simulated full-length unfolded protein. Bars show differences between mean forward MBAR estimates from three replicas per leg. Error bars and numerical labels give *±*1 bootstrap standard error, obtained by independently resampling the replicas within each leg. With three replicas per leg, these errors reflect observed replica variability and do not establish convergence or account for unsampled conformations. Positive values denote destabilization in this cycle. S273F used 40 ns per window.

In the first study, we simulated three linear kinase– substrate complexes containing ATP and two Mg ions (abltide, AblTide and Abl Consensus), with ten 600 ns replicas per complex (18 *µ*s). Over 300–600 ns, the substrate with Gly at position +4 (AblTide) contacted fewer kinase residues per frame than the Phe and Pro substrates (means 1.14, 2.64 and 2.72, respectively, Fig. 5a). This ordering agrees with steady-state kinetics of the Abl kinase domain, in which AblTide had a six-fold higher Michaelis constant than Abl Consensus (35 versus 6 *µ*M) at similar turnover numbers (29).

In the second study, we examined how grafting the same three substrate motifs into the cyclic cystine-knot scaffold MCoTI-II affects groove engagement. Starting from 150 Boltz-2 (30) co-folded models screened by geometric criteria, we built six complexes crossing two binding poses (loop1-in and loop6-in) with the three sequences in the engaged loop while keeping the non-engaged loop as Abl Consensus, and ran ten 300 ns replicas per complex (18 *µ*s). The Abl Consensus pair therefore compared identical molecules in two poses. Over 100–300 ns, all 30 loop1-in replicas remained bound in every frame (mean 32.8 contact pairs for Abl Consensus versus 22.8 in loop6-in, Holm-adjusted permutation *P* = 0.0079), yet their acceptor Tyr5 stayed 5.7–7.3 Å from the catalytic base Asp363 (Fig. 5b). In contrast, loop6-in allowed sequence-dependent catalytic approach: AblTide stayed bound in all ten replicas and held Tyr33 at 3.12 Å from Asp363, closer than Abl Consensus (6.46 Å, *P* = 0.036, 7.3% unbound frames) and abltide (14.38 Å, *P* = 0.015, 10.6% unbound frames). This agrees with the NMR structure of a doubly grafted MCoTI-II, in which loop 1 is more constrained than loop 6 and none of its constrained conformations match the kinase-bound form, whereas loop 6 may adopt it (31). Both studies’ trajectories and conditions can be traced through the DAG.

### Simulation plans evaluate hypotheses proposed by an AI scientist

We then used hypotheses generated by the biomedical AI scientist Biomni (4) as the starting point for a simulation study, corresponding to the AI scientist input in Fig. 1a. PNG-1, the *Neurospora crassa* orthologue of peptide:*N*-glycanase, lacks enzymatic activity but is essential for cell polarity (32). The temperature-sensitive *un-7* allele, which carries S273F together with I390V, cannot grow at the restrictive temperature, and the *png-1* allele carries S279L (33). S273F alone also stops growth at the restrictive temperature (S.H., unpublished). We therefore asked Biomni for its structural cause, and it proposed five hypotheses on disrupted core packing, loss of a buried hydrogen bond, reduced thermal stability, folding destabilization and altered conformations (Fig. 5c). A researcher asked a coding agent to test all five hypotheses in the saved dialogue with MDClaw, substituting other methods where needed, within about 24 h on parallel GPUs. The agent translated them into runs MDClaw supports, using fixed-temperature runs at 400 and 450 K instead of temperature-ramp annealing, 36 ordinary-MD trajectories at 298/310 K, and hybrid-topology free-energy perturbation (FEP) instead of thermodynamic integration. It trimmed the protein to fit the time limit and computed observables lacking an MDClaw tool.

The FEP calculations (Fig. 5d) gave folding free-energy changes of +5.22 *±*0.11, +0.42 *±*0.24, +4.14*±* 0.50 and +0.05*±* 0.03 kcal mol^*−*1^ for S273F, S273A, S279L and I390V. The researcher approved a further 24 h after a promising first S273F estimate, and the agent extended its foldedleg windows to reduce the error. S273F was destabilizing within the range Biomni predicted (+3 to +8), S273A was small and I390V, the second un-7 substitution, near neutral, which partially supports H4. This demonstrates hypothesis testing, from planning to execution across all five hypotheses within the approved time, rather than confirmation of all five.

## Discussion

Setup is the hardest part of automating MD and must be verified at many levels. Consistent with this, most MDCrow failures were on preparation checks (Fig. 3g). We built MD-DataBench from the MDDB, whose deposited simulations come from studies with carefully prepared systems, and developing MDClaw against it enabled AI scientists and agents to use MD autonomously. MDDataBench grades outputs, so the value of the DAG emerged from the process audit, not from the benchmark.

A central idea of MDClaw is that the same DAG both constrains execution and records the process. The ablation separates these roles. The validating tools account for most of the gain in repeated success, whereas the DAG determines whether the process behind a successful run can be checked. Without it, some successful runs had skipped or disconnected stages. For research questions without a reference, this record is what allows a result to be checked.

This approach complements MD agents built on tool use, multi-agent planning and pipeline generation (13–15). Like CHARMM-GUI (11), MDClaw encodes expert protocols, here as validating command-line tools for coding agents. Ai-iDA records calculations in computational materials science as a graph (34), and ScienceClaw records the lineage of artifacts produced by autonomous research agents as a DAG (35).

Two limitations remain. First, the DAG makes every step inspectable, but it is not tamper-proof, because an agent with write access could alter its records, and it covers only calculations run through the tools. A deployment can keep the records where the agent cannot write. The DAG also cannot judge whether a force field, starting model or collective variable suits a scientific question. Second, reliable execution does not overcome the sampling limits of MD. In the PNG-1 study, only one of the five hypotheses was partially supported within the time limit. MD surrogate models and AI structure predictors (30, 36), whose distributions do not always match equilibrium populations (37), could be added as skills or tools to extend sampling.

Beyond working with AI agents on hypotheses, MDClaw can serve the simulation community as a way to acquire data. Carefully set up simulations run by agents could supply AI models with dynamics data that are still scarce, such as antibody ensembles, metalloproteins and flexible complexes, with recorded provenance. The same workflow could compare force fields across many systems and optimize them. MDClaw should also become faster and simpler to use. By making agent-run simulations checkable, recoverable and reusable, MDClaw connects automated hypothesis generation to physics-based validation.

## Methods

### MDClaw architecture and software environment

MD-Claw comprises 14 domain skill documents and a command-line interface, mdclaw, that exposes 100 tools. The skills are concise Markdown files that specify standard protocols for structure preparation, equilibration, production, trajectory analysis, reporting, cluster job management, and alchemical free-energy or enhanced-sampling workflows. Enhanced sampling covers simulated solute tempering (SST2) (38), well-tempered metadynamics (39) and weighted-ensemble sampling (40), whose rounds of walker segments are recorded as ordinary production nodes. Freeenergy workflows cover hybrid-topology relative free energies of point mutations (41) and absolute binding free energies with Boresch restraints (42), analysed with MBAR (43). Example applications are shown in Supplementary Fig. 2. The tools retrieve coordinates from the PDB (44) and sanitize them with PDBFixer, rebuild side chains with H-Packer (45), select chains and ligands, assign titratable residue states with PDB2PQR and PROPKA (46, 47), parameterize proteins, nucleic acids, small molecules, glycans and lipids with AmberTools (10) (for example, OL15 and OL3 for DNA and RNA (48, 49), GAFF (50), GLYCAM06 (51) and Lipid21 (52)), build solvated or membrane-embedded systems with Packmol and Packmol-Memgen (53, 54), and run minimization, equilibration, production and analysis with OpenMM (9), MDTraj (55) and MDAnalysis (56, 57). Ligand partial charges are assigned by default using the OpenFF NAGL graph neural network model, which predicts AM1-BCC-like charges (58). If NAGL is unavailable or charge assignment fails, the tools fall back to AM1-BCC (59). The sum of the partial charges is checked against the ligand’s formal charge.

Unless a task specifies otherwise, proteins use ff19SB (60) in OPC water (61), with a 4 fs time step enabled by hydrogen mass repartitioning (62), a Langevin middle integrator (63) and particle-mesh Ewald electrostatics (64) with a 1.0 nm cutoff. Coding agents (Codex, Claude Code, Open-Code and pi) read the skills through their native skill mechanisms and call mdclaw from the shell. All tools also run without an agent. Analysis is deliberately left open. Language models know libraries such as MDAnalysis and MD-Traj well, so MDClaw provides few analysis tools. Agents write their own scripts to analyze trajectories from any perspective and register the results in analysis nodes of the DAG (`register_analysis_result`), which keeps analyses as flexible as the question requires. The software is distributed as an Apptainer container and a Pixi environment with pinned dependencies.

### Programmatic input validation and error codes

Each tool validates its inputs before any computation. During preparation, checks verify molecular selection (for example, rejecting crystallization artifacts or ligands whose bond connectivity does not match the coordinates), force-field coverage of all residues, net charge, and solvent and counterion parameters. Before a simulation stage, tools verify that thermodynamic parameters match the request, that the parent stage type permits the transition and has completed, and that inputs agree with the recorded ancestry. When a check fails, the tool stops and returns a structured error code with a specific remediation, such as the command needed to parameterize a non-covalently bound amino acid as a ligand. Because the checks are in the tools, they apply whether or not the agent has read the skills.

### Workflow provenance graph

Each workflow stage is stored on disk as a node in a directed acyclic graph (DAG). Node metadata, written by the tool rather than the agent, record the tool, input and output paths, parameters, a hash of the simulated system, execution state, error code on failure and parent node. Failed and superseded nodes are kept. Retries and alternatives are added as new nodes from a suitable ancestor. Because the DAG lives outside the language model’s context, another researcher or agent session can inspect it and resume or branch from a compatible completed node under the same checks. Two tools operate on the DAG: `generate_md_report` drafts a Methods section with software citations from the executed parameters, and `export_mddb` packages metadata, coordinates and trajectories for MDDB deposition, leaving fields that cannot be derived from the record blank.

### MDDataBench task curation

Each task asks an agent to reproduce a reference simulation deposited in the MDDB (27). From its eight repositories (35,602 deposited simulation datasets), we retained all-atom classical MD of a single PDB entry with a stated temperature, a supported explicit water model, an accessible topology, precomputed RMSF and radius-of-gyration analyses, enough frames for at least ten analysis windows and an open licence. In our initial survey, 4,036 datasets in the three largest repositories were eligible. We selected 128 development tasks stratified across nine system classes (Supplementary Table 1).

Prompts and grading contracts were generated by a script from each entry’s metadata and reference files. Each prompt gives only the boundary conditions: PDB identifier, chains and residue ranges, force field and water model, temperature and ensemble, and minimum production length. If the reference modified the PDB coordinates (for example, by removing a fusion tag), the prompt states this in one sentence. Prompts never reveal the MDDB accession. Reference files are retrieved only at grading time and are not redistributed.

### MDDataBench grading protocol

Submissions are graded only from their artifacts: the prepared topology and OpenMM System, minimized coordinates, state logs and production trajectory. Agent-written summaries are not used. Twelve preparation checks compare the system with the reference: chain count and sequences, heavy-atom and hydrogen counts per residue (with exemptions for histidine tautomers, metal-coordinating cysteines and catalytic dyads), elemental composition, disulfides, force-field coverage, net charge, water model, bilayer presence for membrane targets, and a finite energy that decreases on minimization. Eight MD checks test the configured ensemble and temperature, trajectory length, effective elapsed time from water self-diffusion, kinetic temperature, solvent density, Spearman correlation of the RMSF profile, RMSF magnitude and mean radius of gyration. Acceptance bands were derived from reference analysis windows of 1–2.5 ns and widened by four window standard deviations. Calibration details are given in the task contracts. A run passed when every scored check passed. For 5X0T, radius of gyration was diagnostic only and was excluded from the pass criterion. The checks do not establish equilibrium convergence or biological correctness.

### Test set selection

Test systems were selected without using agent outcomes. Reference trajectories required a mean backbone RMSD of at most 4.0 Å over the final third. Sequence similarity was measured as k-mer containment, |*K*(*a*) *∩K*(*b*) | */* min(| *K*(*a*) |, |*K*(*b*) |), with 3-mers for protein and 6-mers for nucleic-acid chains; immunoglobulin chains were compared through their V domain only. A candidate was excluded if the whole system, or any chain of at least 30 residues, reached a containment of 0.70 against a development task or a structure mentioned in the MDClaw code, or if it shared a PDB identifier with either. Category quotas and sequence diversity determined the final set of 31 systems, including the membrane protein 7DTD (human Na_v_1.1 in DPPC).

### Benchmark execution

The three main conditions ran on the RIKYU supercomputer at RIKEN R-CCS (one NVIDIA B200 GPU per MD job) with the pi coding agent and four locally hosted open-weight models with extended reasoning: Kimi-K3, DeepSeek-V4.1-flash, GLM-5.3-flash and Qwen3.8-27B. All conditions used identical prompts: (1) *MDClaw tools + skills*, (2) *MDClaw tools only*, with skills withheld, and (3) *container only*, with the same scientific packages (OpenMM, AmberTools, Packmol-Memgen and PDBFixer) but without MDClaw. Each task was run three times per model and condition. An agent had 30 min to prepare the system and submit Slurm jobs, and each GPU job was limited to 20 min. A filesystem sandbox isolated runs and hid reference data and grading contracts. Network access remained available for structure and package retrieval.

### Statistical analysis

The primary metric, `pass^3`, is the fraction of tasks passed in all three runs. We also report pass@1, the fraction of passing runs, with Wilson 95% confidence intervals. Timeouts, missing artifacts, crashes and validation failures counted as failures. Runs terminated by faults outside the agent’s control (LLM-gateway errors, stopped dispatchers, killed agent processes and a faulty GPU on one compute node) were re-executed, and their original records are kept. The DAG ablation (below) was compared with its control by a two-sided sign test over matched model–system pairs, without adjustment for dependence among models on the same system.

### Harness optimization and comparison

Harness development began with 98 tasks and later included 30 additional tasks selected under the same rules. The 98-task set was evaluated with Kimi-K3, GLM-5.3-flash and Qwen3.6-35B. The 30-task set used Kimi-K3, DeepSeek-V4.1-flash, GLM-5.3-flash and Qwen3.8-27B. Each model–task combination used three conditions and three repeats, giving 2,646 and 1,080 recorded runs, respectively. The task prompts required at least 1 ns of production MD for 105 tasks and 2.5 ns for 23 tasks. Agents examined recorded failures and proposed changes, which a human specialist reviewed before revising the tools and skills. These repeated development rounds changed the harness, not the language-model weights.

The pre-optimization version was the release with which MDDataBench-driven development began; it had already been refined on earlier benchmarks. The pre-optimization and optimized harness versions were evaluated on the same 31 test tasks with identical models, prompts and settings. Each version used its own container and skills, so the comparison evaluates the complete distributed harness. The container-only baseline was not repeated.

### No-DAG ablation

The ablation ran the optimized MDClaw tools with the DAG layer disabled by an environment variable, in the same container image. Tools that create, inspect or branch nodes were hidden, together with the study, round-based sampling and free-energy workflows that depend on them. Each remaining tool takes explicit file paths and writes fixed, documented file names into a directory the agent chooses. No conditions, such as temperature, box size, disulfide bonds or ligand charge, are inherited from earlier stages, and tool guidance that refers to nodes is removed. The chemistry code and input checks are those of the standard harness. Each call also writes a record of its tool, arguments, exit status, and input and output files with SHA-256 hashes, which the agent does not need to read. Prompts were those of the tools-only condition, except that the agent wrote a submission file naming its final production directory. For grading, the per-call records were used to reconstruct the stages leading to that production and to assemble the files the grader reads from a DAG, so the checks were unchanged. The control was the tools-only condition on the same 30 non-membrane systems, four models and three runs, and the two were compared with the sign test described above.

The process audit followed each scored production back to its inputs. A run had a process error when equilibration or production did not start from the output of the preceding stage, when a structure entering a stage was not the recorded output of an earlier stage or had changed after it was written, or when a stage was missing. With the DAG, the same questions were answered from node parents, node status and the system hashes recorded by each node. The DAG records do not hash the starting state of production, which the tool reads from its parent node. Failure episodes were counted per run from the transcript and Slurm accounting: an episode begins at a tool result reporting failure, or at an MD job that did not complete, and ends when the same stage later succeeds or the run ends.

### Token usage and recorded workflow progress

Token usage per run (Fig. 3c) is the sum of prompt and output tokens over all model calls, including cached input and reasoning tokens, for all 1,116 runs of the three main conditions. For Fig. 3g, we read the grader’s record of each unsuccessful run. A run had no gradable production when it had no score or when every weighted check failed for the same reason, such as a missing submission file or no completed production node; otherwise each failed weighted check was counted.

### MDCrow baseline evaluation

We ran the public, unmodified MDCrow (13) on the 31 test tasks with DeepSeek-V4.1-flash and GLM-5.3-flash (three runs each), using the same prompts and following its published configuration. Because MDCrow runs simulations within the agent process, each run was a single 90 min GPU job. Aggregate metrics exclude the unsupported membrane task (180 runs over 30 systems). Compatibility fixes in the LangChain parsing layer made model output-format errors recoverable. MDCrow’s tools and prompts were not changed. An OpenMM-level hook captured MDCrow’s outputs for grading without modifying them.

### Abl1 substrate and cyclotide simulations

The investigators designed both studies and posed the questions and comparisons. For the linear Abl1–substrate complexes (Y.W.), an agent ran Boltz-2 co-folding (five samples per substrate), the active-state screen, the rebuilding of ATP and two Mg ions from PDB 1ATP, system building, equilibration and 30 production runs through MDClaw. The investigator inspected the selected models and fixed one common Tyr4 starting geometry (from PDB 1IR3) for the three systems. For the Abl1– cyclotide complexes (Y.Z.), an agent cut 150 Boltz-2 candidates to six systems with executable geometric gates and carried the work from sequence to trajectory in one DAG (126 production nodes, 186 Slurm jobs), deriving the mutated complexes by branching from one template. Manual intervention was needed to pin histidine states and disulfides, to transplant ATP and two Mg ions from PDB 1ATP, to apply four patches to MDClaw for the head-to-tail cyclic scaf-fold, and to keep the analysis outside the DAG. The analyses of both studies used scripts outside the DAG. For the linear Abl1–substrate complexes (abltide, AblTide and Abl Consensus, with ATP and two Mg ions), we ran ten 600 ns replicas per substrate with ff19SB/OPC at 300 K following a 0.5 ns common Tyr4-geometry restraint stage. The +4 contact count is the number of kinase residues within 0.45 nm of the substrate residue. Fig. 5a uses per-replica averages over 300–600 ns, compared with 150–300 ns as a check. For the Abl1–cyclotide complexes, 150 Boltz-2 (30) co-folded models (50 per sequence with physical steering potentials) were screened for active kinase geometry, groove engagement (acceptor Tyr OH within 0.6 nm of Asp363), three cystine-knot disulfides (0.18–0.23 nm) and head-to-tail closure. After transplanting ATP and two Mg ions from PDB 1ATP, six complexes (loop1-in and loop6-in crossed with abltide, AblTide and Abl Consensus in the engaged loop, with the non-engaged loop held as Abl Consensus) were derived from the shared Abl Consensus template, solvated with ff19SB/OPC in 0.15 M NaCl, minimized in two stages, equilibrated over four stages (3 ns) and simulated in ten 300 ns replicas per complex (18 *µ*s total). Head-to-tail backbone cyclization, exclusion of cyclic chains from terminal capping, mutation to glycine and ancestor lookup of ligand chemistry were executed from a local MDClaw worktree branch. Observables were averaged over bound frames (at least one residue pair within 0.45 nm) in 100–300 ns after translation-only anchor imaging, and compared across replicas by two-sided permutation tests (20,000 shuffles) with Holm adjustment across all 159 single-variable comparisons.

### PNG-1 hypothesis testing and free-energy analysis

A researcher gave a saved Biomni dialogue to a coding agent and asked it to test all five hypotheses with MDClaw, using parallel GPUs within approximately 24 h and substituting available methods where MDClaw lacked a proposed one. From then on the agent worked automatically through MD-Claw, from the study plan to the analyses. It wrote a plan that mapped each hypothesis to simulations and observables with criteria for support, refutation and inconclusive results, prepared and equilibrated the systems, ran the simulations, and analyzed them, including the MBAR free-energy estimates and the difference between the two legs of each cycle. It used constructs of residues 47–368 and 47–412 because the full-length system (345,000 atoms) did not fit the time limit. Observables without an MDClaw tool (solvent-accessible area, hydrogen bonds, secondary structure, principal components, water counts and Lennard-Jones energies) were computed with mdtraj and OpenMM on the MDClaw trajectories and registered in the DAG. The starting model was entry Q7SI01 of the AlphaFold Protein Structure Database (65), with a structural zinc placed at the centroid of the C150, C153, C182 and C185 sulfurs (treated as thiolates), and ff19SB/OPC. The plan replaced temperature-ramp annealing with 100 ns NVT runs at 400 and 450 K (24 trajectories), alongside 36 ordinary MD trajectories at 298 and 310 K.

Because the first S273F estimate looked promising, the researcher approved a further 24 h, which the agent used to extend the S273F folded-leg windows from 10 to 40 ns to reduce the error. Folding free-energy changes at 310 K were computed as the difference between alchemical transformations in the folded protein and a tripeptide reference (41), with positive values denoting destabilization. Soft-core potentials were used for steric interactions (66). Each leg had three replicas, using 21 or 31 windows and 5–40 ns per window, and was analyzed with MBAR (43) after automated equilibration detection (67). Fig. 5d shows the difference of replica means. Errors are bootstrap standard errors over replicas and do not establish convergence. S279L was a related buried-serine substitution, I390V a predicted nearneutral comparator, and S273A was included to distinguish proposed hydrogen-bond and side-chain-volume effects.

### Use of large language models

Four open-weight models were evaluated on MDDataBench as described above, two of them also with MDCrow. In the application studies, coding agents ran the Abl1 substrate and cyclotide simulations with MDClaw. For PNG-1, Biomni (4) generated the hypotheses and an initial validation protocol through its public web interface. Claude Code (Anthropic) with Claude Opus 5.5 then planned and executed the simulations and their analyses through MDClaw. Claude Code, Cursor and Codex were used to inspect benchmark failure logs, implement software and analysis code, prepare figures, and draft and revise the manuscript. The authors specified the design, grading criteria and task selection, and reviewed and take responsibility for all content.

## Data availability

MDDataBench task definitions, prompts, scoring contracts and calibration records are available at https://github.com/matsunagalab/MDDataBench. Source data for the figures, including per-run benchmark outcomes, token counts, DAG-stage summaries, Abl1 substrate and cyclotide replica-level measurements and PNG-1 free-energy estimates, together with the per-run records of the MDDataBench evaluation (results, scores, agent transcripts and DAG records) and the DAG records, analysis outputs and reduced trajectories of the three application studies, are available in Zenodo (https://doi.org/10.5281/zenodo.24095749). The source records specify calculation versions and input paths where available. Reference simulations were obtained from MDDB (https://mddbr.eu) and are not redistributed. Task definitions record the originating node, accession, retrieval date and licence. Starting structures were obtained from the Protein Data Bank and, for PNG-1, the AlphaFold Protein Structure Database (Q7SI01, version 6).

## Code availability

MDClaw is available under the MIT licence at https://github.com/matsunagalab/mdclaw, and MDDataBench, including the scorer and benchmark runner, at https://github.com/matsunagalab/MDDataBench.

## ACKNOWLEDGEMENTS

We thank the organizers and mentors of the RIKEN AI4S Hackathon 2026, hosted by NVIDIA and the RIKEN Center for Computational Science, where the first applications of MDClaw were developed. Computations were performed on the RIKYU supercomputer at the RIKEN Center for Computational Science.

## Author contributions

Y.M. conceived the study, led the development of MDClaw and MDDataBench, and drafted the manuscript. S.K. proposed constructing the benchmark from deposited molecular dynamics simulation databases. A.E. and Y.H. contributed to the implementation of the MDClaw tools. Y.W. and Y.Z. carried out the Abl1 substrate and cyclotide application studies, respectively. S.K., Y.W. and Y.Z. tested MDClaw on application studies and provided feedback and suggestions that improved the harness. S.H. provided biological context on PNG-1 and contributed to scientific discussions of the simulation results. A.S. supervised the application studies. Y.M., S.K., Y.W., Y.Z., S.H. and A.S. contributed to revising the manuscript.

