## Supplementary Information for "An auditable harness for biomolecular dynamics simulations by AI agents"

### Contents

Supplementary Fig. 1. Methods draft, references and MDDB deposition bundle assembled from a recorded workflow.
Supplementary Fig. 2. Enhanced-sampling and free-energy workflows run with MDClaw.
Supplementary Table 1. MDDataBench tasks.

**a** Recorded workflow (5OD1 job)

- `source_001`
  PDB 5OD1 (mmCIF), SHA-256 recorded
- `prep_001`
  Chain A 1–94, ligand 9RQ (−2);
  PDB2PQR + PROPKA, pH 7.0
- `solv_001`
  TIP3P, cubic, 15 Å buffer,
  0.15 M NaCl; 39,115 atoms
- `topo_001`
  ff99SB-ILDN; GAFF2 + NAGL charges;
  PME, 1.0 nm cutoff; HMR
- `min_001`
  ≤5,000 steps; solute heavy-atom
  restraints, 100 kJ mol$^{-1}$ nm$^{-2}$
- `eq_001`
  298 K, 1 bar; 0.1 ns NVT +
  0.25 ns NPT; 4 fs
- `prod_001`
  1.5 ns NPT; frames every 10 ps

generate_md_report reads these node records
and writes report.json and references.bib.

**b** References selected from recorded evidence

| BibTeX key | Role | Evidence: node, file, field |
|---|---|---|
| `Eastman2024OpenMM8` | software | `topo_001 system.xml` `/System/@openmmVersion` |
| `Zhang2019LFMiddle` | method | `prod_001 integrator.xml` `/Integrator/@type` |
| `Chow1995MCBarostat,` `Aqvist2004MCBarostat` | method | `prod_001 runtime_system.xml` `Force[@type='MonteCarloBarostat']` |
| `Hopkins2015HMR` | method | `topo_001 node.json` `/metadata/hmr` |
| `Jorgensen1983TIP3P` | parameters | `solv_001 node.json` `/metadata/water_model` |
| `Wang2004GAFF,` `Wang2006Antechamber` | parameters, software | `topo_001 node.json` `…/small_molecule_forcefield` |
| `Wang2025AshGCWorkingPaper` | method | `topo_001 node.json` `…/ligand_charge_assignment` |
| `Olsson2011PROPKA3,` `Dolinsky2004PDB2PQR` | method, software | `prep_001 node.json` `/metadata/protonation_baseline_method` |
| `SchottVerdugo2019PackmolMemgen` | software | `solv_001 solvated.pdb` `REMARK (Packmol-Memgen)` |

18 references were selected, each tied to a node, file and field (9 shown). Reported as unresolved: the protein force-field paper (the ff99SB-ILDN mapping is not automated) and the constraint solver. Versions without a dedicated paper: openmmforcefields 0.16.0, gaff-2.11, OpenFF NAGL model openff-gnn-am1bcc-1.0.0.

**c** Methods paragraph drafted by a coding agent from the report

The protein–ligand complex was taken from PDB entry 5OD1 (chain A, residues 1–94, with ligand 9RQ). Protonation states at pH 7.0 were assigned with PDB2PQR and PROPKA [Dolinsky2004PDB2PQR, Dolinsky2007PDB2PQR, Jurrus2018APBSPDB2PQR, Olsson2011PROPKA3, Sondergaard2011PROPKA], giving HID at His35, His61 and His65. The ligand carried a net charge of −2 and was parameterized with GAFF2 [Wang2004GAFF] through Antechamber and AmberTools [Wang2006Antechamber, Case2023AmberTools], with partial charges from a graph neural network AM1-BCC model [Wang2025AshGCWorkingPaper]. The protein was modelled with the ff99SB-ILDN force field and water with TIP3P [Jorgensen1983TIP3P]. The complex was solvated in a cubic box with a 15 Å buffer and 0.15 M NaCl (33 $Na^+$ and 31 $Cl^-$; 39,115 atoms) using Packmol-Memgen [SchottVerdugo2019PackmolMemgen, Martinez2009Packmol]. Simulations were run with OpenMM 8.5.1 [Eastman2024OpenMM8] using particle-mesh Ewald electrostatics with a 1.0 nm cutoff, constrained bonds to hydrogen and hydrogen mass repartitioning [Hopkins2015HMR], which allowed a 4 fs time step. After up to 5,000 steps of energy minimization with 100 kJ mol$^{-1}$ nm$^{-2}$ restraints on solute heavy atoms, the system was equilibrated with solute heavy-atom restraints for 0.1 ns at constant volume and 0.25 ns at 1 bar, at 298 K, with the Langevin middle integrator (friction 1 ps$^{-1}$) [Zhang2019LFMiddle] and a Monte Carlo barostat [Chow1995MCBarostat, Aqvist2004MCBarostat]. Production ran for 1.5 ns under the same conditions without restraints, with coordinates saved every 10 ps.

Checklist returned with the draft

Add the ff99SB-ILDN reference (not supplied by the report).

Ligand protonation used Dimorphite-DL (pH 7.0); no reference was supplied.

State the constraint solver; the record does not identify SETTLE, SHAKE or CCMA.

Cite openmmforcefields 0.16.0, gaff-2.11 and the NAGL model by version.

One 1.5 ns run; no random seed recorded; no claim of convergence.

**d** MDDB deposition bundle (export_mddb; not uploaded)

```
mddb/
├── project/
│   ├── inputs.yaml
│   └── md_001/
│       ├── system.pdb          1,518 atoms; water and counter-ions removed
│       └── trajectory.dcd      150 frames, stride 1; no imaging or fitting
├── manifest.json               source SHA-256, atom indices, checks
├── report.json                 same report as in b and c
└── references.bib              BibTeX for the selected keys
```

Checks before export: PDB and DCD atom and frame counts, atom ordering, first-frame coordinates, YAML round trip and the MDDB inputs schema.

project/inputs.yaml (excerpt)


```
name: 5OD1 protein-ligand complex (MDClaw demo)
license: CC BY 4.0
type: trajectory
input_structure_filepath: md_001/system.pdb
mds:
- name: main:prod_001
  input_trajectory_filepaths:
  - md_001/trajectory.dcd
  program: OpenMM
  version: 8.5.1
  temp: 298.0
  timestep: 4.0
  ensemble: NPT
  ff: [amber/ff99SBildn.xml, amber/tip3p_standard.xml]
  wat: tip3p
  method: Unbiased molecular dynamics
  metadditions:
    mdclaw: {source files and SHA-256, node history, ...}
```


**Supplementary Fig. 1. Methods draft, references and MDDB deposition bundle assembled from a recorded workflow.** The job is a protein–ligand complex (PDB 5OD1, ligand 9RQ) from a development benchmark run (GLM-5.3-flash, tools plus skills). **a**, Recorded DAG of the job with selected recorded settings of each node. `generate_md_report` reads these node records and writes `report.json` and `references.bib`. **b**, Nine of the 18 selected references. Each reference is tied to the node, file and field that show the method was used. Methods that the report cannot map to a reference are listed as unresolved, and versioned resources without a dedicated paper are listed separately. **c**, Methods paragraph drafted by a coding agent (Claude Code) from `report.json` and `references.bib`, following the MDClaw reporting skill, which cites only the supplied BibTeX keys and lists missing facts and references in a checklist instead of inventing them. The draft is shown unedited. **d**, Offline MDDB deposition bundle written by `export_mddb`. The trajectory is reduced to the solute, and the manifest records the source files with SHA-256 hashes and the checks run before export. The bundle was not uploaded.

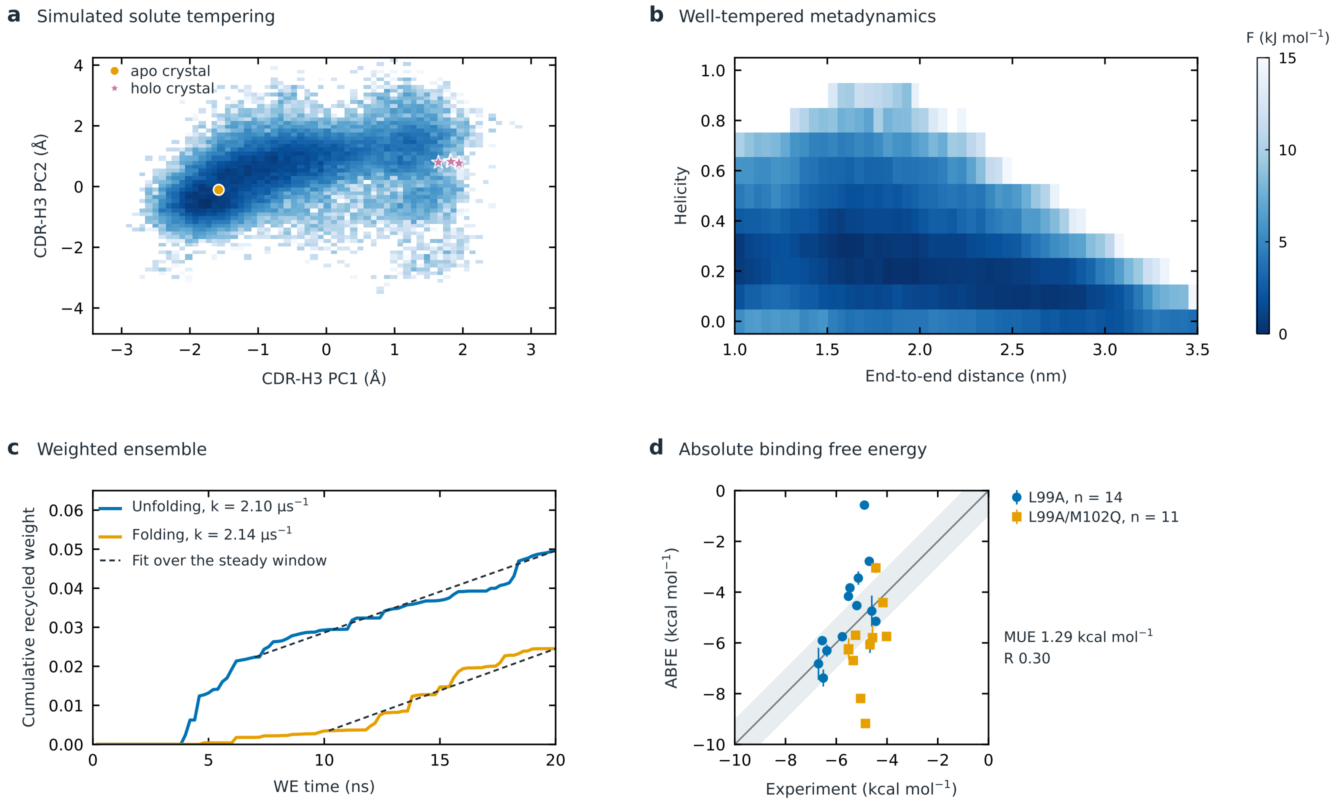


**Supplementary Fig. 2. Enhanced-sampling and free-energy workflows run with MDClaw.** Each calculation was planned and recorded as nodes of an MDClaw DAG. Unless stated otherwise, systems used ff19SB and OPC with hydrogen mass repartitioning and a 4 fs time step. The colour scale of the free-energy maps in **a** and **b** is common. **a**, Simulated solute tempering (SST2) (1) of the CDR-H3 loop of an anti-CTLA-4 nanobody, started from the apo crystal structure (PDB 5E03), with only CDR-H3 as the solute. Free energy at 300 K (eight walkers $\times$ 300 ns, MBAR weights) on the first two principal components of CDR-H3 backbone coordinates after superposition of the framework. Circle, apo crystal; stars, the four copies of the holo crystal structure (PDB 5E5M). Six of eight walkers reached the holo-like CDR-H3 torsion pattern at 300 K, whereas three 300 ns runs of plain MD from the same start did not. **b**, Well-tempered metadynamics (2) of capped deca-alanine on the end-to-end distance (four walkers $\times$ 100 ns sharing one bias, bias factor 8, 300 K). Frames were reweighted with the final bias. Helicity is the fraction of residues with backbone dihedrals in the $\alpha$-helical region. **c**, Weighted-ensemble simulation (3) of the folding and unfolding of chignolin CLN025 at 340 K (100 rounds of 0.2 ns with about 150 walkers each per direction). The slope of the cumulative weight recycled into the target state over the steady window gives the rate constant: 2.10 (unfolding) and 2.14 $\mu s^{-1}$ (folding), compared with 3.13 and 1.74 $\mu s^{-1}$ from four 1 $\mu$s unbiased simulations with the same force field. **d**, Absolute binding free energies of 25 ligands to T4 lysozyme L99A and L99A/M102Q by double decoupling with Boresch restraints (4) (GAFF2 with AM1-BCC charges for ligands, three replicas, 5 ns per window), against isothermal titration calorimetry. The band marks $\pm$1 kcal mol$^{-1}$.

**Supplementary Table 1. MDDataBench tasks.** All 159 tasks of MDDataBench v0.7.1: 128 development tasks used while building MDClaw and 31 test tasks used for the evaluation in the main text, grouped by system class. Each task reproduces one reference simulation deposited in the MDDB; the line under each class gives the MDDB collection that provided the references, with its publication where available, and the force field, water model and simulation program of the references. Task, number in the task identifier; PDB, starting structure; MDDB, accession of the reference simulation; System, the simulated chains, ligands and bilayer; $T$, reference temperature. Prompts never reveal the MDDB accession.

| Task | PDB | MDDB | System | $T$ (K) |
|---|---|---|---|---|
| **Development tasks (128)** | | | | |
| *Soluble proteins (ATLAS)* (29) | | | ATLAS (5), repository bsc · CHARMM36m; TIP3P; GROMACS | |
| 064 | 16PK | A02K9 | 3-Phosphoglycerate kinase (*T. brucei*) | 300 |
| 065 | 1A62 | A02KA | Rho termination factor RNA-binding domain (*E. coli*) | 300 |
| 066 | 1AB1 | A02KB | Crambin, Ser22/Ile25 form (*C. hispanica*) | 300 |
| 067 | 1AF7 | A02KC | Chemotaxis receptor methyltransferase CheR (*S. typhimurium*) | 300 |
| 068 | 1AIL | A02KE | NS1 RNA-binding domain (influenza A) | 300 |
| 069 | 1AOL | A02KF | Envelope glycoprotein gp70 receptor-binding domain (Friend MLV) | 300 |
| 070 | 1BGF | A02KL | STAT4 N-terminal domain (mouse) | 300 |
| 071 | 1BQ8 | A02KN | Rubredoxin (*P. furiosus*) | 300 |
| 072 | 1BXY | A02KQ | Ribosomal protein L30 (*T. thermophilus*) | 300 |
| 073 | 1C1K | A02KT | Gene 59 helicase assembly protein (phage T4) | 300 |
| 074 | 1C52 | A02KU | Cytochrome c552 (*T. thermophilus*) | 300 |
| 075 | 1CPQ | A02KZ | Cytochrome c′ (*R. capsulatus*) | 300 |
| 076 | 1CTF | A02L0 | Ribosomal protein L7/L12 C-terminal domain (*E. coli*) | 300 |
| 077 | 1DOW | A02LF | $\beta$-Catenin–$\alpha$-catenin chimera (mouse) | 300 |
| 078 | 1DPT | A02LG | D-Dopachrome tautomerase (human) | 300 |
| 079 | 1EWF | A02LV | Bactericidal/permeability-increasing protein (human) | 300 |
| 080 | 1EZ3 | A02LW | Syntaxin-1A N-terminal domain (rat) | 300 |
| 081 | 1FCZ | A02M2 | Retinoic acid receptor $\gamma$ ligand-binding domain (human) | 300 |
| 082 | 1FD3 | A02M3 | $\beta$-Defensin 2 (human) | 300 |
| 083 | 1FK5 | A02M5 | Nonspecific lipid-transfer protein (maize) | 300 |
| 084 | 1FM4 | A02M6 | Major birch pollen allergen Bet v 1-L (birch) | 300 |
| 085 | 1G38 | A02MF | DNA methyltransferase M.TaqI (*T. aquaticus*) | 300 |
| 086 | 1G6G | A02MH | Rad53 FHA1 domain (yeast) | 300 |
| 087 | 1GQV | A02MP | Eosinophil-derived neurotoxin (human) | 300 |
| 121 | 2O6X | A0311 | Procathepsin L1 (*F. hepatica*) | 300 |
| 122 | 1J8R | A02NS | PapG pyelonephritic adhesin receptor-binding domain (*E. coli*) | 300 |
| 123 | 3LAE | A03DG | UPF0053 protein HI0107 (*H. influenzae*) | 300 |
| 124 | 2CWY | A02WZ | Hypothetical protein TTHA0068 (*T. thermophilus*) | 300 |
| 125 | 2P54 | A031Y | PPAR$\alpha$ ligand-binding domain (human) | 300 |
| *Soluble proteins (MoDEL)* (18) | | | MoDEL (6), repository mmb · Amber Parm99; TIP3P; Amber | |
| 088 | 12CA | A0001 | Carbonic anhydrase II (human) | 300 |
| 089 | 1A1W | A000G | FADD death effector domain (human) | 300 |
| 090 | 1AA3 | A000N | RecA C-terminal domain (*E. coli*) | 300 |
| 091 | 1AG4 | A0015 | Spherulin 3a (*P. polycephalum*) | 300 |
| 092 | 1AH9 | A001D | Translation initiation factor IF1 (*E. coli*) | 300 |
| 093 | 1ANX | A000H | Annexin V (human) | 300 |
| 094 | 1APO | A000C | EGF-like module of coagulation factor X (bovine) | 300 |
| 095 | 1ARD | A000J | ADR1 zinc-finger domain (yeast) | 300 |
| 096 | 1AY7 | A001N | Ribonuclease Sa (*S. aureofaciens*) with barstar (*B. amyloliquefaciens*) | 300 |
| 097 | 1DFW | A00BS | Lung surfactant protein B fragment | 300 |
| 098 | 1TBA | A011R | TBP (yeast) with TAF1 N-terminal domain (*D. melanogaster*) | 300 |
| 099 | 1VYC | A014L | Bucain three-finger toxin (*B. candidus*) | 300 |
| 100 | 2AXF | A002J | HLA-B*3508–$\beta$2-microglobulin with a BZLF1 10-mer peptide (human) | 300 |
| 126 | 1PWT | A00W4 | $\alpha$-Spectrin SH3 domain (chicken) | 300 |
| 127 | 1J7A | A00O3 | Ferredoxin I (Nostoc sp. PCC 7119) | 300 |
| 128 | 1AJJ | A001O | LDL receptor ligand-binding module 5 (human) | 300 |
| 129 | 2CRO | A008D | Cro repressor (phage 434) | 300 |
| 130 | 1C2A | A009E | Bowman–Birk trypsin inhibitor (barley) | 300 |
| *Protein–ligand complexes* (13) | | | LIGATE, repository cin · Amber ff99SB-ILDN; TIP3P; GROMACS | |
| 036 | 1CEB | A0004 | Plasminogen kringle 1 (human) with tranexamic acid (AMH) | 298 |
| 037 | 1G74 | A0006 | Adipocyte lipid-binding protein mutant (mouse) with oleic acid (OLA) | 298 |
| 039 | 3IKD | A000F | Pin1 peptidyl-prolyl isomerase (human) with inhibitor J9Z | 298 |
| 040 | 3N2U | A002A | MMP-12 catalytic domain (human) with inhibitor D3X | 298 |
| 041 | 4ERF | A000I | MDM2 N-terminal domain (human) with AM-8553 (0R3) | 298 |
| 042 | 4MN3 | A000J | CBX7 chromodomain (human) with a methyllysine peptidomimetic (LIG) | 298 |
| 043 | 5OD1 | A000O | Engineered metalloesterase MID1sc10 with phosphonate (9RQ) | 298 |
| 044 | 5OH3 | A000P | Cereblon isoform 4 (*M. gryphiswaldense*) with ethosuximide (9V2) | 298 |
| 045 | 6J3O | A002C | PCAF (KAT2B) bromodomain (human) with compound B4L | 298 |
| 102 | 4MRE | A00DC | CD44 hyaluronan-binding domain (mouse) with ligand 2C9 | 298 |
| 103 | 5YJ8 | A000Y | TDRD3 Tudor domain (human) with ligand 8W9 | 298 |
| 104 | 3S0E | A00AR | Odorant-binding protein 14 (honey bee) with eugenol (EOL) | 298 |
| 105 | 6MJ7 | A0002 | p62/SQSTM1 ZZ domain (human) with arginine (ARG) | 298 |

Supplementary Table 1 (continued)

| Task | PDB | MDDB | System | $T$ (K) |
|---|---|---|---|---|
| *Protein–protein complexes* (10) DynaRepo, repository inr · CHARMM36m; TIP3P; GROMACS | | | | |
| 025 | 1AKJ | A000B | HLA-A*0201–$\beta$2-microglobulin–HIV RT peptide with CD8$\alpha\alpha$ (human) | 310 |
| 026 | 1AVA | A000C | Barley $\alpha$-amylase 2 with $\alpha$-amylase/subtilisin inhibitor (barley) | 310 |
| 027 | 1B6C | A000F | FKBP12 with TGF-$\beta$ type I receptor cytoplasmic domain (human) | 310 |
| 028 | 1DFJ | A000H | Ribonuclease A (bovine) with ribonuclease inhibitor (porcine) | 310 |
| 029 | 1E3U | A000J | OXA-10 class D $\beta$-lactamase homodimer (*P. aeruginosa*) | 310 |
| 030 | 1FFW | A000R | CheY with the CheY-binding domain of CheA (*E. coli*) | 310 |
| 107 | 1LJ2 | A001D | Rotavirus NSP3 RNA-binding domain with an eIF4G peptide | 310 |
| 108 | 4MQV | A007Y | RPA32 C-terminal domain with a SMARCAL1 peptide (human) | 310 |
| 109 | 5SXP | A009Z | $\beta$-PIX SH3 domain with an Itch peptide (human) | 310 |
| 110 | 3K1R | A005C | Harmonin with SANS (USH1G) C-terminal fragment (human) | 310 |
| *Nanobodies* (8) Nanobodies, repository mmb · Amber ff99SB-ILDN; TIP3P; GROMACS | | | | |
| 031 | 1KXV | A02JY | VHH cAb10 against pancreatic $\alpha$-amylase (dromedary) | 300 |
| 032 | 1XFP | A02JZ | VHH cAb-Lys3 CDR2 germline revertant (dromedary) | 300 |
| 033 | 3TPK | A057R | Oligomer-specific VHH KW1 (camelid) | 300 |
| 034 | 4KFZ | A057Y | Anti-LMO2 VH single domain (human) | 300 |
| 035 | 6GWN | A0594 | VHH-2g-42 against PAI-1 (alpaca) | 300 |
| 115 | 8IJZ | A05CE | Anti-VEGF nanobody | 300 |
| 116 | 6NI2 | A059A | Nanobody 32 (llama) | 300 |
| 117 | 7VNE | A05BH | Nanobody n3113.1 | 300 |
| *Antibody–antigen and other protein complexes* (13) Dynabench, repository inr · CHARMM36m; TIP3P; NAMD | | | | |
| 015 | 1AHW | A00KU | Tissue factor extracellular domain (human) with Fab 5G9 (mouse) | 300 |
| 016 | 1AY7 | A00MF | Ribonuclease Sa (*S. aureofaciens*) with barstar (*B. amyloliquefaciens*) | 300 |
| 017 | 1DQJ | A00KV | Hen egg-white lysozyme with Fab HyHEL-63 (mouse) | 300 |
| 018 | 1MLC | A00KW | Hen egg-white lysozyme with Fab D44.1 (mouse) | 300 |
| 019 | 2DD8 | A00KY | SARS-CoV spike receptor-binding domain with neutralizing Fab (human) | 300 |
| 020 | 2VIS | A00L1 | Influenza A hemagglutinin HA1 escape mutant T131I with Fab (mouse) | 300 |
| 021 | 3EOA | A00L4 | LFA-1 (integrin $\alpha$L) I domain with efalizumab Fab (human) | 300 |
| 022 | 3RVW | A00LC | Der p 1 (house dust mite) with Fab 4C1; entry obsolete, superseded by 5VPG | 300 |
| 023 | 3WD5 | A00LF | TNF-$\alpha$ with adalimumab Fab (human) | 300 |
| 024 | 5CBA | A00LO | CXCL13 with Fab 3B4 (human) | 300 |
| 118 | 3MXW | A00LA | Sonic hedgehog with Fab 5E1 | 300 |
| 119 | 5Y9J | A00LX | BAFF (TNFSF13B) with belimumab Fab (human) | 300 |
| 120 | 3L5W | A00L8 | Interleukin-13 with Fab C836 | 300 |
| *Nucleic acids* (18) BIGNASim (7), repository mmb · force field not recorded; TIP3P; Amber | | | | |
| 046 | 1A66 | A01DE | 12-bp DNA duplex from the NFATc1–DNA complex (protein removed) | 300 |
| 047 | 1C7U | A01DF | 20-bp self-complementary DNA duplex from the MEF2A–DNA complex | 300 |
| 048 | 1H9T | A01DH | 19-bp DNA duplex, fadB operator from the FadR complex | 300 |
| 049 | 1IV6 | A01DJ | 13-bp telomeric DNA duplex from the TRF1–DNA complex | 300 |
| 050 | 1J46 | A01DK | 14-bp DNA duplex from the SRY HMG box–DNA complex | 300 |
| 051 | 1KX5 | A017E | Nucleosome core particle, 147-bp DNA with histone octamer (*X. laevis*) | 298 |
| 052 | 1NAJ | A01A6 | 12-bp DNA duplex (Dickerson–Drew dodecamer) | 298 |
| 053 | 1QN5 | A01DO | 14-bp TATA-box DNA duplex from the TBP complex (protein removed) | 300 |
| 054 | 1ZGW | A01DS | 18-bp DNA duplex | 300 |
| 055 | 2HDC | A01DT | 17-bp DNA duplex | 300 |
| 056 | 2L1G | A01DV | 16-bp DNA duplex | 300 |
| 057 | 2LT7 | A01DW | 19-bp DNA duplex | 300 |
| 058 | 2RN1 | A017Z | 16-bp RNA duplex | 300 |
| 059 | 3F27 | A01DZ | 16-bp DNA duplex from the Sox17–DNA complex | 300 |
| 111 | 1JE8 | A01DM | 20-bp self-complementary DNA duplex | 300 |
| 112 | 1HLV | A017B | 21-bp CENP-B box DNA duplex from the CENP-B complex; SPC/E water | 298 |
| 113 | 3JXC | A01E0 | 20-bp self-complementary DNA duplex | 300 |
| 114 | 1JRN | A01DC | Bimolecular G-quadruplex of d(G4T4G4) (Oxytricha telomere) | 300 |
| *Membrane proteins* (15) EBRAINS/MCNS, repository mmb · CHARMM36; TIP3P; GROMACS | | | | |
| 001 | 5YC8 | A01M3 | M2 muscarinic acetylcholine receptor, fusion partner removed (human); DPPC bilayer | 300 |
| 002 | 5ZK3 | A01M7 | M2 muscarinic acetylcholine receptor, fusion partner removed (human); DPPC bilayer | 300 |
| 003 | 5ZK8 | A01M9 | M2 muscarinic acetylcholine receptor, fusion partner removed (human); DPPC bilayer | 300 |
| 004 | 5ZKB | A01MA | M2 muscarinic acetylcholine receptor, fusion partner removed (human); DPPC bilayer | 300 |
| 005 | 6A93 | A01MB | 5-HT2A serotonin receptor, fusion partner removed (human); DPPC bilayer | 300 |
| 006 | 6A94 | A01MC | 5-HT2A serotonin receptor, fusion partner removed (human); DPPC bilayer | 300 |
| 007 | 6GT3 | A023T | A2A adenosine receptor, fusion partner removed (human); DPPC bilayer | 310 |
| 008 | 6I53 | A023K | GABA-A receptor $\alpha1\beta3\gamma2$ pentamer (human); DPPC bilayer | 310 |
| 009 | 6JZH | A024E | A2A adenosine receptor, fusion partner removed (human); DPPC bilayer | 310 |
| 010 | 6KUX | A024G | $\alpha$2A adrenergic receptor (human); DPPC bilayer | 310 |
| 011 | 6KUY | A024H | $\alpha$2A adrenergic receptor (human); DPPC bilayer | 310 |
| 012 | 6ME3 | A023M | MT1 melatonin receptor, fusion partner removed (human); DPPC bilayer | 310 |
| 013 | 6PS2 | A023O | $\beta$2 adrenergic receptor, T4 lysozyme fusion removed (human); DPPC bilayer | 310 |
| 014 | 6ZDV | A023S | A2A adenosine receptor, fusion partner removed (human); DPPC bilayer | 310 |

Supplementary Table 1 (continued)

| Task | PDB | MDDB | System | $T$ (K) |
|---|---|---|---|---|
| 101 | 7DFP | A024X | D2 dopamine receptor, fusion partner removed (human); DPPC bilayer | 310 |
| *Metalloproteins* (4) | | | CV19 (3); LIGATE (1), repository cin, mmb · Amber ff14SB, Amber ff99SB-ILDN; TIP3P; GROMACS | |
| 060 | 4OW0 | MCV1900208 | SARS-CoV papain-like protease with structural zinc | 298 |
| 062 | 6W9C | MCV1900209 | SARS-CoV-2 papain-like protease (nsp3) with structural zinc | 298 |
| 063 | 6WRH | MCV1900210 | SARS-CoV-2 papain-like protease (nsp3) with structural zinc | 298 |
| 106 | 6RNT | A0008 | Ribonuclease T1 (*A. oryzae*) with 2′-AMP (2AM) | 298 |
| **Test tasks (31)** | | | | |
| *Soluble proteins (ATLAS)* (7) | | | ATLAS (5), repository bsc · CHARMM36m; TIP3P; GROMACS | |
| 148 | 1HOE | A02N0 | $\alpha$-Amylase inhibitor tendamistat (*S. tendae*) | 300 |
| 149 | 5N76 | A03TC | CooT nickel chaperone (*R. rubrum*) | 300 |
| 150 | 1KPT | A02OH | KP4 killer toxin (*U. maydis*) | 300 |
| 151 | 2X9O | A035Q | 15,16-Dihydrobiliverdin:ferredoxin oxidoreductase PebA (Synechococcus sp.) | 300 |
| 152 | 2FP1 | A02YH | Secreted chorismate mutase (*M. tuberculosis*) | 300 |
| 153 | 1D4T | A02L6 | SAP SH2 domain (human) | 300 |
| 154 | 2PEF | A032C | Serpin tengpin (*T. tengcongensis*) | 300 |
| *Soluble proteins (MoDEL)* (6) | | | MoDEL (6), repository mmb · Amber Parm99; TIP3P; Amber | |
| 155 | 1QIB | A00XJ | MMP-2 catalytic domain (human) | 300 |
| 156 | 1A5D | A004D | $\gamma$E-Crystallin (rat) | 300 |
| 157 | 1HP9 | A00JB | $\kappa$-Hefutoxin 1 (scorpion) | 300 |
| 158 | 1EDG | A00DP | Endoglucanase A catalytic domain (*C. cellulolyticum*) | 300 |
| 159 | 1CS6 | A00AF | Axonin-1 immunoglobulin domains 1–4 (chicken) | 300 |
| 160 | 1CEX | A008J | Cutinase (*F. solani pisi*) | 300 |
| *Protein–ligand complexes* (4) | | | LIGATE, repository cin · Amber ff99SB-ILDN; TIP3P; GROMACS | |
| 131 | 2PQL | A00BY | D7r4 salivary protein (*A. gambiae*) with tryptamine (TSS) | 298 |
| 132 | 3QXV | A00E1 | Anti-methotrexate CDR-grafted VHH (llama) with methotrexate (MTX) | 298 |
| 133 | 5EQ1 | A00IS | BRPF1 (peregrin) bromodomain (human) with ligand BEA | 298 |
| 134 | 5I7X | A00E2 | BRD9 bromodomain (human) with ligand 67B | 298 |
| *Protein–protein complexes* (4) | | | DynaRepo, repository inr · CHARMM36m; TIP3P; GROMACS | |
| 135 | 2D10 | A002T | Radixin FERM domain (mouse) with an EBP50 peptide | 310 |
| 136 | 4KT5 | A007O | GrlR–GrlA regulator complex (*E. coli*) | 310 |
| 137 | 1I51 | A000Y | Caspase-7 with the XIAP BIR2 domain (human) | 310 |
| 138 | 5GY2 | A00BJ | TLR5–VLRB fusion (zebrafish/hagfish) with flagellin (*B. subtilis*) | 310 |
| *Nanobodies* (3) | | | Nanobodies, repository mmb · Amber ff99SB-ILDN; TIP3P; GROMACS | |
| 143 | 6IR2 | A0598 | mCherry-binding nanobody LaM2 (Bactrian camel) | 300 |
| 144 | 6X05 | A059P | VHH-SAN4 (alpaca) | 300 |
| 145 | 6B73 | A058W | Nanobody (llama) | 300 |
| *Antibody–antigen and other protein complexes* (2) | | | Dynabench, repository inr · CHARMM36m; TIP3P; NAMD | |
| 146 | 5O14 | A00LR | Factor H binding protein variant 1.1 (*N. meningitidis*) with Fab 1A12 (human) | 300 |
| 147 | 5X0T | A00LW | Basigin (human) with Fab 6H8 (mouse) | 300 |
| *Nucleic acids* (4) | | | BIGNASim (7), repository mmb · force field not recorded; TIP3P; Amber | |
| 139 | 2STW | A01DY | 17-bp DNA duplex | 300 |
| 140 | 4F6N | A01E2 | 19-bp DNA duplex with 5-methylcytosines | 300 |
| 141 | 1VTN | A01DQ | 13-bp DNA duplex | 300 |
| 142 | 1A0A | A01DD | 17-bp DNA duplex | 300 |
| *Membrane proteins* (1) | | | EBRAINS/MCNS, repository mmb · CHARMM36; TIP3P; GROMACS | |
| 161 | 7DTD | A0251 | Nav1.1 sodium channel $\alpha$ subunit (human); DPPC bilayer | 310 |
